# Slow cortico-cortical connectivity (2-5Hz) is a new robust signature of conscious states

**DOI:** 10.1101/692970

**Authors:** Pierre Bourdillon, Bertrand Hermann, Marc Guénot, Hélène Bastuji, Jean Isnard, Jean-Rémi King, Jacobo Sitt, Lionel Naccache

## Abstract

While long-range cortico-cortical functional connectivity has been reported by several studies as a necessary condition of conscious state, precise empirical evidence is still scarce. In the present work we provide such a direct and conclusive evidence in a set of three experiments. In the two first experiments intracranial-EEG was recorded during four distinct states in the same individuals: conscious wakefulness (CW), rapid-eye-movement sleep (REM), stable periods of slow-wave sleep (SWS) and deep propofol anaesthesia (PA). We discovered that long-range FC, computed by the weighted Symbolic-Mutual-Information (wSMI) in the 2-5Hz frequency band was a specific marker of conscious states that could discriminate CW and REM from SWS and PA. In the third experiment, we generalized this original finding on a large cohort of brain-injured patients by revealing that wSMI in the 2-5 Hz range was also able to accurately discriminate patients in the vegetative state (or unresponsive wakefulness syndrome) from patients in the minimally conscious state. Taken together the present results suggest that 2-5Hz FC is a new and robust signature of conscious states.

## Introduction

Identifying a robust neural signature of conscious states constitutes a prominent scientific and medical challenge. While several theoretical and empirical works put forward the importance of cortico-cortical connectivity for conscious states (Amico et al., 2017; Boly et al., 2013; Crone et al., 2014; Dehaene and Changeux, 2011; Dehaene and Naccache, 2001), this property remains highly debated regarding : (i) the long-range (Laureys and Schiff, 2012) versus short-range type of functional connectivity (Lamme, 2006) (ii) the relevant frequencies involved in this cortico-cortical connectivity, with contradicting proposals ranging from ultra-slow (<0.01Hz) (Barttfeld et al., 2015), to slow (0.5-4Hz) (He et al., 2008; He and Raichle, 2009), or even faster theta, alpha and gamma-band rhythms (Schiff et al., 2014), and (iii) the general value of this property when comparing various conscious and unconscious states.

In order to address these important and unsolved questions, we explored cortico-cortical connectivity on direct intracranial EEG (iEEG) recordings performed in the same subjects across various conscious and unconscious states. More precisely, we recorded iEEG in 12 drug-resistant epileptic patients undergoing a stereoelectroencephalography (SEEG) for pre-surgical evaluation with large and brain-scale implantations. Each patient was recorded in two states that are associated with conscious experience (conscious wakefulness (CW) and rapid-eye movement sleep (REM) that is typically associated with conscious dreaming), as well as in two unconscious states: very stable periods of slow-wave sleep (SWS) that are usually free of conscious dreaming (Siclari et al., 2017), and deep propofol anaesthesia (PA).

We estimated cortico-cortical connectivity by computing the weighted symbolic mutual information (wSMI) in various frequency bands. We previously conceived wSMI and used this measure to distinguish conscious and minimally conscious (MCS) patients from vegetative state/unresponsive wakefulness syndrome (VS/UWS) patients (King et al., 2013). wSMI evaluates the extent to which two EEG signals present non-random linear or non-linear joint fluctuations. This method probes qualitative (‘symbolic’) patterns of information sharing in the signal (see Supplemental Experimental Procedures). The symbolic transformation depends on the number of the symbols and on their temporal separation (sampling of EEG values each τ) enabling to probe various frequency bands. The wSMI algorithm was recently used to reveal an increase of cortico-cortical after restoration of consciousness by vagus nerve stimulation in one patient in the VS/UWS (Corazzol et al., 2017). In these two studies, wSMI computed in the 4-10Hz (τ=32ms) distinguished MCS from VS/UWS states, while wSMI computed in higher frequencies failed to do so.

We conducted two consecutive SEEG studies: in study I, five patients were recorded during 4 nights in three stages (CW, REM, SWS), whereas in study II, 7 additional patients were recorded during short periods of 10 minutes in four stages (CW, REM, SWS and PA). Finally, we conducted a third study using high-density scalp EEG on 145 patients suffering from disorder of consciousness (MCS and VS/UWS) (Naccache, 2018), in order to determine if our findings would discriminate between these two states.

## Results

In coherence with our original study, wSMI computed in the 4-10Hz succeeded to distinguish CW from SWS in 8 out of the 12 recorded patients: mean wSMI 4-10Hz computed across all bipolar contacts combinations was larger during CW than during SWS (t-test p<0.001 for each patient) in each of the 8 patients, including all 5 patients of study I with long-lasting recordings. However, three of the remaining four patients showed a reverse pattern (larger mean wSMI 4-10Hz values during SWS than during CW; p<0.001) and one of them did not show significant difference between these two states. More surprisingly, this measure of connectivity failed to discriminate robustly other conscious (CW and REM) from unconscious states (selected very stable and sustained samples of SWS and PA). Indeed, only 3 out of 7 patients showed a significant larger mean wSMI 4-10Hz for CW than PA, whereas the four other patients show the reverse pattern with higher mean wSMI 4-10Hz for PA. Similarly, wSMI 4-10Hz was higher during REM than during SWS in only 5 out of 12 patients, while the 7 remaining patients showed an opposite significant pattern. Finally, only 2 out of 7 patients showed larger wSMI 4-10Hz in REM than in PA, while the five remaining ones also showed the reverse pattern (see Figure 1a and Figure S2b).

**Figure 1.**
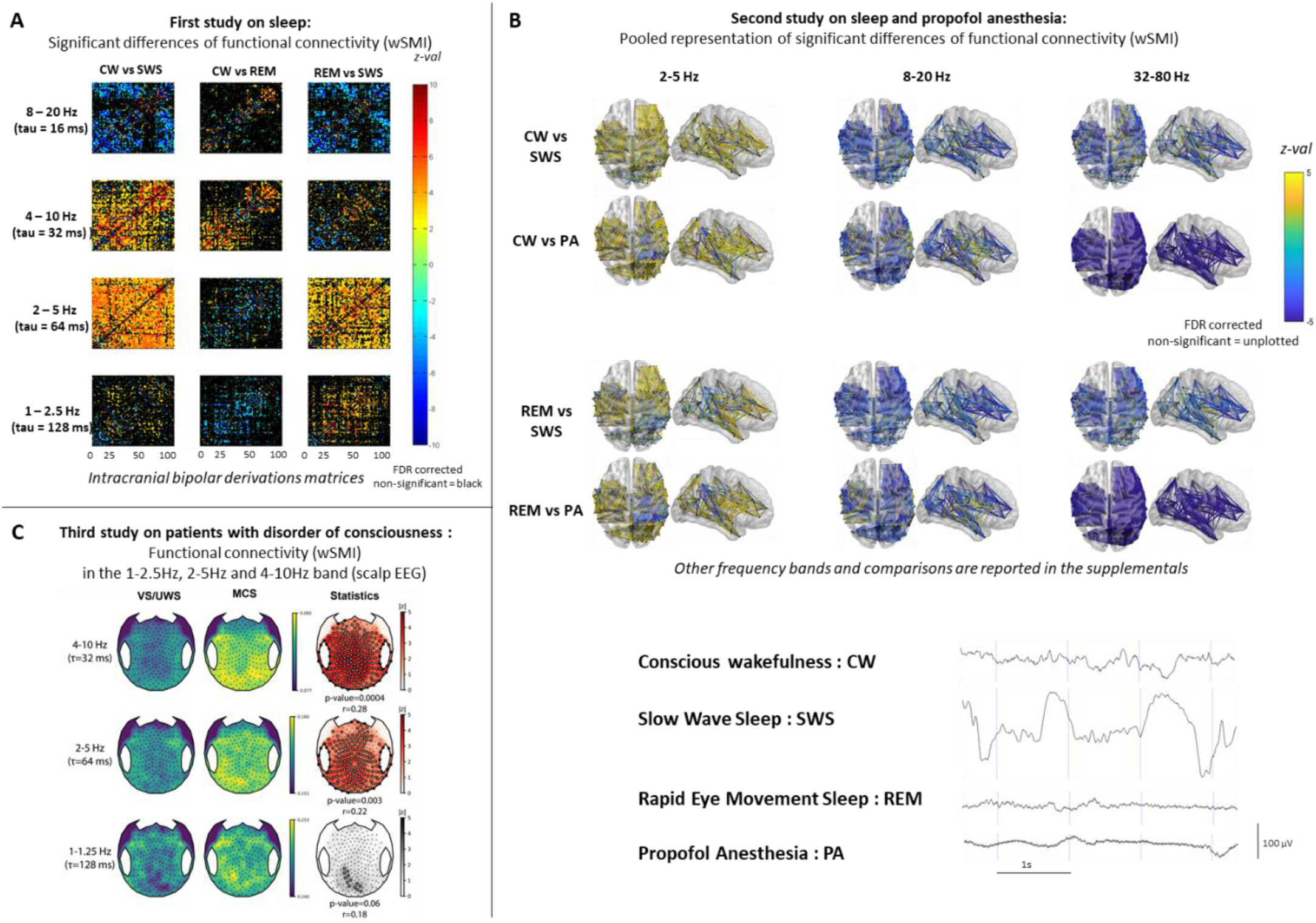
Electrophysiological signature of consciousness and specificity of propofol induced loss of consciousness. (a.) Example of raw iEEG signal of each of the studied stages in the two first experiment. A: Matrices (bipolar derivations x bipolar derivations) of statistical comparison of the mean wSMI values between stages. Color bar corresponds to the z-values. Non-significant results are plotted in black. B: Anatomical representation of significant differences (z-values) of wSMI between stages in 2-5 Hz, 8-20 Hz and 32-80 Hz frequency bands. z-values of non-significant differences (after FDR correction) are not plotted. Data from the 7 patients are represented in a common MNI space. C: Scalp EEG topographic representation of wSMI values of VS/UWS and MCS patients (left columns) and results from the permutation cluster-based statistics (right column). Absolute z-values are plotted with a red colorscale when a significant cluster was found and in grey otherwise with the corresponding p-value and effect size r of the cluster. Electrodes belonging to clusters are highlighted by white circles. On the bottom, example of raw iEEG signal of each of the studied stages in the two first experiment. CW = conscious wakefulness; SWS = N3 Slow Wave Sleep; REM = Rapide Eye Movement Sleep; PA = propofol induced general anaesthesia; UWS = unresponsive wakefulness state; VS/UWS = vegetative state/unresponsive wakefulness syndrome; MCS = minimally conscious state;; wSMI = weighted Symbolic Mutual Information

We then discovered that the wSMI calculated on slower frequencies (wSMI 2-5Hz; τ=64ms) actually succeeded much better to discriminate robustly the two conscious states from the two unconscious ones: 10 out of 12 patients showed significant larger mean values during CW than during SWS, while the remaining two patients (who both had short recordings) did not show significant differences across these two states. The very same result was found when comparing REM to SWS. A mirror observation was made in the 8-20 Hz (τ=16ms) and in the 32-80Hz (τ=4ms) where mean wSMI was significantly larger (*p*<*0*.*001*) in the SWS as compared to CW and REM in all patients (Figure 1 b). Concerning PA, all seven patients had a larger wSMI 2-5Hz during CW than PA, and during REM than PA (see Figure 1b and S2c). As a control test, we checked that wSMI computed in a slower frequency band (1-2.5Hz; τ=128ms) did not perform well to discriminate conscious from unconscious states (Fig 1a and S1c).

We also noticed that PA, unlike SWS, was associated with a massive, diffuse and systematic increase of functional connectivity in high frequencies (32-80Hz (τ=2ms), see Figure 1b) as compared to the three other states. For each comparison and each patient, more than 90% of the pairs of electrodes showed this effect. These findings are congruent with the increase of gamma band coupling recently described in non-human primates on ECoG during experimental anaesthesia with propofol and with ketamine (Bola et al., 2017).

In order to probe the generalization of our new measure of conscious state to other alterations of consciousness, we finally tested if wSMI 2-5Hz could discriminate brain activity of patients in the MCS from patients in the VS/UWS.Using 167 five minutes resting state high-density EEG recordings acquired from 145 distinct patients (N=68 VS/UWS and N=77 MCS), we first showed that wSMI in the 4-10 Hz was able to discriminate VS/UWS from MCS (mean scalp wSMI AUC=0.66 CI_95_[0.57-0.74], p=0.0005, effect size r=0.27 with a significant cluster encompassing almost all scalp electrodes, p=0.0004), replicating the original findings of King et al. where wSMI was computed during an active paradigm task (King et al., 2013). Crucially, we then found that wSMI in the 2-5 Hz range was also able to accurately discriminate VS/UWS from MCS (mean scalp wSMI AUC=0.62 [0.53-0.70], p=0.01, r=0.20, with a significant cluster encompassing almost all the scalp, p=0.004), while wSMI in the 1-1.25 Hz range was not (mean scalp wSMI AUC=0.54 [0.45-0.62], p=0.42, r=0.06 with no significant cluster, p=0.11) (see Figure 1c). Similar results were found when computing wSMI during the local-global active paradigm as originally described (data not shown).

## Discussion

In the present iEEG study, we first showed that long-range cortico-cortical connectivity measured with the wSMI computed in the theta-alpha band (4-10Hz), - that was previously shown to be higher for patients in the MCS from those in the VS/UWS -, was larger during CW than during SWS. However, and unexpectedly, this measure failed to discriminate both REM from SWS, and REM and CW from PA. In other terms, FC in the 4-10Hz does not seem to be a general signature of conscious states. We then discovered that FC computed in a slower delta-theta band (2-5Hz) actually succeeded much better to discriminate correctly conscious states (CW and REM) from the two unconscious states investigated here (SWS and PA). This measure was found larger in CW and in REM as compared to SWS and to PA in the vast majority of patients. Notably, none of the 12 recorded patients showed a reverse pattern of connectivity between conscious and unconscious states in this delta-theta band. Finally, we could generalize the validity of this new measure in another population of patients and with high-density scalp EEG recordings: wSMI computed in this 2-5Hz band was significantly larger in the MCS than in the VS/UWS. As a control condition, we showed that wSMI computed in a slower band (1-2.5Hz) failed to discriminate conscious from unconscious states, both in iEEG and in scalp-EEG. Taken together, our results suggest that cortico-cortical connectivity in the delta-theta band could correspond to a general neural signature of consciousness that is valid across a large variety of impairments of consciousness. In addition to enriching cognitive neuroscience of consciousness, the present data and its proposed interpretation may impact clinical practice with a new neurophysiological signature of conscious states.

This proposal is compatible with a core element of the Global Workspace (GW) theory according to which conscious states would correspond to the serial chaining of ∼200-300ms discrete states (Koch et al., 2016; Milz et al., 2016), each associated with a P3b signature (Naccache, Ph. Trans. 2018). Indeed, this hypothesis leads to a 3.3-5Hz frequency range of coherent and synchronized brain-scale patterns that is ideally captured by the wSMI 2-5Hz. Moreover, our proposal seems also very coherent with electrophysiological and fMRI data that highlighted the importance of slow cortical potentials (SCP) in cognition and consciousness (He et al., 2008). He and Raichle listed some explanatory arguments in favor of such a link between SCP (0.1-4Hz) and conscious states. In particular, they underlined the fact that “long-range cortico-cortical connections terminate preferentially in superficial layers, and thus contribute significantly to SCP” (He and Raichle, 2009).

In the light of these results and of previous reports, we propose a two-frequency hypothesis of cortical connectivity that introduces a distinction between signatures of conscious state and signatures of conscious access.

In two previous SEEG studies, we identified brain-scale increases of connectivity in the alpha-beta band during conscious access to written words (Gaillard et al., 2009), and in the alpha band during conscious access to the regularity auditory series of tones (Bekinschtein et al., 2009; Sitt et al., 2014). We then identified, - in high-density scalp EEG recordings -, a sharp difference of long-range connectivity within the theta-alpha band between controls in a CW state and patients in the MCS on the one hand, and patients in the VS/UWS on the other hand (King et al., 2013; Sitt et al., 2014). This progressive slow-down of frequency windows of connectivity when moving from conscious access contrasts, to conscious states contrasts tentatively suggests a mechanistic difference between conscious access and conscious state. We propose that conscious states would require a lasting level of long-range functional connectivity within a delta-theta band enabling a sustained level of coherence and information integration between the distant neural processors contributing to the GW, whereas conscious access would occur by the mean of sudden and transient (within the second time-range) increases of connectivity between a specified cortical network and the GW. In other terms, while brain-scale increases in FC in the theta-beta band would reflect sudden conscious accesses, FC in the slower delta-theta band would capture the ongoing activity of a functional conscious GW. This last measure seems more specific than the proposed signature of conscious access, but the reasons of this lack of specificity require additional studies.

Finally, we will also discuss our findings related to increase of FC in the gamma-band during unconscious states. Recently, loss of consciousness has also been related to hyper-correlated gamma-band activity in anesthetized macaques and sleeping humans through intracranial EEG recordings (Bola et al., 2017). We replicated this finding in the present study and generalized it for the first time to anesthetized human subjects. The enhanced inter-dependence of gamma-band activity during alteration of consciousness may reflect suppression of information transfer (Chialvo, 2010) consecutive to the well-established decrease of the complexity of electrophysiological signals during unconscious states (Alonso et al., 2014; Casali et al., 2013; Krzemiński et al., 2017; Solovey et al., 2015; Tajima et al., 2015). Gamma activity is indeed frequently considered as a macro-scale reflection of a bursty activity pattern observed during sleep and anesthesia (Lewis et al., 2012; Nir et al., 2011; Vizuete et al., 2014). Massive positive correlations may results from bursty aspecific activities occurring at multiple brain locations. Another interpretation could be that the phase of low-frequency oscillations (Lakatos et al., 2008), changes in cross-frequency interactions between delta and gamma could possibly drive the observed enhanced functional connectivity (Bola et al., 2017).

## Materials and Methods

### First and second experiment

The two first experiments of this study used a very similar approach. We therefore report their respective experimental procedures in a single section.

#### Patients

All patients had a drug-resistant epilepsy requiring phase II investigation by stereo-electroencephalography. They benefited from this procedure in in the department Neurosurgery of Lyon university hospital (Hospital for neurology and neurosurgery Pierre Wertheimer, Lyon University, France). Data, including intracranial-EEG (iEEG) during wakefulness, sleep and general anaesthesia, were collected anonymously and no change the routine management of patients was needed. More specifically the acquisition of intracranial-EEG (iEEG) under general anaesthesia was performed during the surgical implantation and did not modify the duration of the procedure. This study was approved by the local ethics comity (Comité Consultatifs de Protection des Personnes se Prêtant à des Recherches Biomédicales. Authorization No. 22236S).

None of the 5 patients of the first experiment had any sleep disorder history. No modifications of anti-epileptic drugs occur during or between the recordings. Patient 1 did not receive any antiepileptic drug; Patient 2 received Levetiracetam 2000mg & Lamotrigin 800mg; Patient 3 received Levetiracetam 1000mg & Lamotrigin 800mg; Patient 4 received Carbamazepin 600mg & Pregabalin 75mg; Patient 5 received Carbamazepin 800mg & Valproate 500mg.

None of the 7 patients of the second experiment had any sleep disorder history. No modifications of anti-epileptic drugs occur during or between the recordings. Patient 1 received Lamotrigin 800mg & Perampanel 12mg; Patient 2 received Levetiracetam 1000mg & Lamotrigin 800mg; Patient 3 received Pregabalin 75mg & Carbamazepin 800mg; Patient 4 received Valproate 500mg & Carbamazepin 600mg; patient 5 received Levetiracetam 2000mg; Patient 6 received Carbamazepin 1200mg & Zonizamide 500mg; Patient 7 received Lamotrigin 800mg.

#### Stereo-electroencephalography

Stereo-electroencephalography (SEEG) was performed under general anaesthesia using a frame based Talairach methodology (Guenot et al., 2001). Electrodes (Microdeep®) were manufactured by Dixi (Dixi Medical, 4, chemin de Palente, BP 889, 25025 Besancon, France). Dimensions of each contact are 2 mm in length, 0.8 mm in diameter and the intercontact spacing is 1.5mm. Each electrode has from 5 to 15 contacts. Signal was recorded by a Micromed® system with frequency sampling of 256Hz.

SEEG was performed under general anaesthesia using Target-Controlled Infusion (TCI) with propofol (2,6 diisopropylphénol) (Debailleul et al., 2004). Deepness of anaesthesia was controlled by measuring the Bispectral-Index (BIS) (Hans et al., 2000).

For the second experiment, the iEEG was recorded (10 min) during the final part of the procedure, during the per-operative imaging control, so before the decrease of anaesthesia and during a perfectly stable level of propofol anaesthesia (PA).

#### Electrodes localisation

All patient had a postoperative MRI in the 24h following the SEEG. All electrodes were manually localized and their location were put into a the MNI space by using MRIcro (Chris Rorden’s MRIcro, Copyright © 1999-2005) (Rorden and Brett, 2000) and SPM 12 (Copyright © 1991,1994-2018 FIL).

Tridimensional representation (see Figure S1 a.) and matching with functional connectivity matrices were computed by using BrainNet Matlab toolbox (v 1.61, released 20171031) (Xia et al., 2013).

#### Sleep staging

Based on American Academy of Sleep Medicine (AASM) recommendation of 2007 (Silber et al., 2007) and adaptation to iEEG (Magnin et al., 2004) sleep staging were performed and then controlled by a sleep expert (HB). Periods of 10 min of time, comparable to those obtained under propofol general anaesthesia, were then selected in stable periods of conscious wakefulness (CW), N3 slow wave sleep (NREM) and rapid eyes movement sleep (REM). Concerning the N3 slow wave sleep, only periods of stable slow wave activities were selected as we wanted conscious content, as dreaming, to be avoided. Dreaming experiences, so associated with form of consciousness, during N3 slow wave sleep have indeed been associated to fragmentations of the slow activities during this stage (Siclari et al., 2017).

#### Data pre-processing

All iEEG data underwent an artefact rejection and a visual analysis using Anywave software (IDDN.FR.001.110001.000.S.P.2014.000.31230 AMU and INSERM) (Colombet et al., 2015). No channel had to be excluded. All paroxistic or pathological epileptic EEG abnormalities were excluded from the selected recording. The 10 min periods of each stage (wakefulness, N3 slow Wave sleep, rapid eyes movement sleep and propofol induced general anaesthesia) was divided into 10s periods of time. A re-referencement of electrode contacts to a bipolar montage with their nearest neighbour on the same physical electrode. Specifically before functional connectivity analysis, non-contiguous electrode contacts were excluded to avoid massive biases.

#### Power spectrum analysis

The power spectrum analysis of each frequency band (δ: [0.5 – 4 Hz[; θ: [4 – 8 Hz[; α: [8 – 12 Hz[; β: [12 – 30 Hz[; γ [30 – 70 Hz]) was performed with Matlab® using the Fieldtrip toolbox (Copyright © 2008-2018, Donders Institute for Brain, Cognition and Behaviour, Radboud University, The Netherlands) (Oostenveld et al., 2011). A multitaper methods and a path of time of 0.5s was used. Each frequency band and each stage was normalized by computing the sum of power in a frequency band reported to the power on all frequency bands of the spectrum sum.

#### Functional connectivity analysis

Functional connectivity was estimated by computing the weighted symbolic mutual information index (wSMI) previously described by King at al. (King et al., 2013) as a relevant measure to discriminate between different states of consciousness. Conversely to phase or amplitude correlation measures, this index, derived from the permutation entropy analysis, can detect non-linear coupling between pair of bipolar derivation. First, signal was reduced into a limited set of discrete 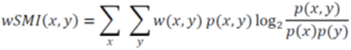 symbols, made of groups of sub-vectors, including a certain number of points (k = 3, so 6 existing patterns) sampled with a particular temporal interval. This temporal interval, τ, determined the frequency range for which the index become sensible. Every sub-vector corresponds to a particular symbol, assigned according to different reciprocal patterns that the three points can assume(Corazzol et al., 2017). Here the frequency bands investigated were 32-80 Hz (τ = 4 ms), 16-40 Hz (τ = 8 ms), 8-20 Hz (τ = 16 ms), 4-10 Hz (τ = 32 ms), 2-5 Hz (τ = 64 ms) and 1-2.5 Hz (τ = 128 ms). The symbols’ probability density and counting their mutual occurrence over two time-series the index was determined prior to compute an estimate of the coupling between each pair of electrodes (Corazzol et al., 2017; King et al., 2013): (x and y being the pairs of symbols of the two time series).

#### Statistical analysis

All statistical analysis was performed with Matlab© R2017a v9.2.0.556344 and the stats toolbox (Copyright © 1984-2017, The MathWorks©, Natick, Massachusetts, USA). A Welch’s test without hypothesis of equal variances was used to perform the statistical comparisons on the normalized power spectrum between the different stages across the 5 frequency bands. A non-parametric two-tail Mann-Whitney U-test was used for comparisons between wSMI values of each state of consciousness. The area under the curve (AUC) of the receiver operating characteristic curve (ROC) was then reported. Note that AUC is related to the U statistic of the Mann-Whitney U test: 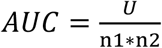 (where n1 and n2 are the size of both groups). Control for multiple comparisons was made by using the False Discovery Rate (FDR) (Benjamini and Hochberg, 1995; Glickman et al., 2014; Hochberg and Benjamini, 1990). The 0.05 threshold of significance was adapted at each test according to this adjustment with the FDR.

Comparison of the different mean wSMI for each frequency band between the stages for each patient was performed by using a student t-test.

### Third experiment

#### Population

Five minutes resting state EEG were acquired from a convenience sample of patients suffering from disorders of consciousness (DoC) assessed in la Pitié-Salpêtrière university hospital (Paris, France) from February 2014 to July 2018. Patients were assessed by trained physicians using the dedicated Coma Recovery Scale revised (Giacino et al., 2002; Kalmar and Giacino, 2005). This specialized scale not only quantifies behavior responses to a set of predetermined and hierarchical items in six different domains (auditory, visual, motor, oro-motor, communication and arousal), but also determines the consciousness state of the patient according to some key behaviors elicited by the passation of the scale. Patients were thus diagnosed as being in a vegetative state/unresponsive wakefulness syndrome (VS/UWS) or in a minimally conscious state (MCS). A total of 203 recordings were acquired from 167 unique DoC patients. After the automated preprocessing pipeline (see below), 167 valid recordings acquired from 145 unique patients were amenable for analysis. The population, 68 VS/UWS and 77 MCS patients, was typical of DoC patients, with 87 males and 54 females (sex-ratio=1.61), median age of 47.4 IQR[30.4-62.6] years. Etiologies were anoxia in 37%, traumatic brain injury in 24%), stroke in 5% and other causes in 34%. Median delay from injury was 58 [31.0-184.0] days. Consent was obtained from the patient’s relative. The protocol conformed to the Declaration of Helsinki, to the French regulations, and was approved by the local ethic committee (*Comité de Protection des Personnes; CPP n° 2013-A01385-40*) Ile de France 1 (Paris, France).

#### EEG acquisition and preprocessing

Five minutes resting state scalp EEG were acquired in a quiet room while the patients received no particular instruction. Recordings were done using a NetAmps 300 Amplifier (Electrical Geodesics, Eugene, Oregon) with a high-density sponge-based 256 channels HydroCel Geodesic Sensor Net (Electrical Geodesics) referenced to the vertex at 250 Hz sampling frequency. Impedances were checked before the beginning of the recording and set below 100 kΩ.

An automatized and hierarchical preprocessing workflow written in Python, C, bash shell scripts and based on open source technologies, including the software MNE (Gramfort et al., 2014), was used for artefact removal and quality assessment (Engemann et al., 2018; Engemann and Gramfort, 2015; Sitt et al., 2014). Shortly, this previously described pipeline followed the subsequent steps: 1) Filtering: raw data were band-pass filtered (0.5 Hz 6^th^-order Butterworth high-pass filter and 45 Hz 8^th^-order Butterworth low-pass filter) with 50 Hz and 100 Hz notch filters. 2) Epoching: filtered data were cut into 800 ms epochs with a 550 to 850 ms random jitter in-between (these timings were chose to reproduce the original wSMI analysis in which wSMI was computed on the 800 ms baseline periode of an active auditory oddball paradigm (King et al., 2013)). 3) Bad channels and bad epochs removal: channels that exceeded a 100 μv peak-to-peak amplitude in more than 50% of the epochs were rejected. Channels that exceeded a z-score of 4 across all the channels mean variance were rejected. This step was repeated two times. Epochs that exceeded a 100 μv peak-to-peak amplitude in more than 10% of the channels were rejected. Channels that exceeded a z-score of 4 across all the channels mean variance (filtered with a high pass of 25 Hz) were rejected. This step was repeated two times. The remaining epochs were digitally transformed to an average reference. Rejected channels were interpolated. Finally, EEG were deemed to pass this preprocessing step if at least 70% of the channels and at least 30% of the epochs were kept.

#### Functional connectivity analysis

As previously described, wSMI was computed on the 800 ms epochs using a k=3 kernel in the different frequency bands, respectively: tau=32 ms (4-10 Hz), tau=64 ms (2-5 Hz) and tau=128 ms (1-1.25 Hz), yielding a single value of wSMI in each frequency band for each pair of scalp electrodes (224 ×(224-1)/2=24976) per epochs and per patient. A single value for each electrode was then obtained by computing the median value of the connectivity between this electrode and every other scalp electrodes, resulting in a measure related to the degree of a network in graph theory analysis. These median values were then averaged over time using the trimmed mean 80%, a robust estimator of central tendency (Wilcox and Rousselet, 2018). A single two-dimensional topography of the 224 scalp electrodes was thus characterizing each patient. Furthermore, the mean wSMI over the scalp was also used as a measure of the overall magnitude of functional connectivity in each frequency band.

#### Statistical analysis

We compared the wSMI in each frequency band between VS/UWS and MCS, both at the level of the mean wSMI over the scalp (wSMI_mean_) and at the topography level. Populations were compared using the non-parametric Mann-Whitney U test. The discriminative power of wSMI_mean_ to distinguish VS/UWS from MCS patients was assessed using the area under the ROC curve (AUC) and its bootstrapped 95% confidence interval computed using 10000 iterations. Note that AUC is related to the U statistic of the Mann-Whitney U test:

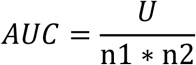

Where n1 and n2 are the size of both groups.

For the topography analysis, we used a cluster-based permutation procedure (Maris et al., 2007; Maris and Oostenveld, 2007) to control for multiple comparisons at sensor level. The first step of this procedure consisted of the comparison of the wSMI values at each 224 sensor, using the same statistical test as previously described. Spatial clusters of z-statistic corresponding to type I error of 5% were then constructed using the neighboring matrix of electrodes. Each resulting clusters were assigned a value, known as the cluster mass, which corresponds to the sum of the z-statistic of the individual components of the cluster. Random permutation (N=10000) of the patient’s label were then used to create a surrogate distribution of the cluster masses under the null hypothesis. One can then compute the probability of each cluster constructed with the original data to be observed under the null hypothesis. In addition to p-values, Effect sizes were reported using the measure *r*:

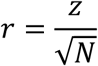

Where *z* is the z-statistic of the Mann-Whitney U test and *N* the size of the population. This effect size measure was computed both at the level of the wSMI_mean_, but also at the level of the cluster (effect size computed on the mean over electrodes belonging to the cluster).

## Author Contributions

PB and LN proposed the concept study, the working hypotheses and supervised the experimental protocol; MG and PB performed surgery and implanted the electrodes; PB performed the iEEG recordings under propofol and JI during the other stages; PB and LN performed the pre-processing and the analysis the analyses; PB and MG performed the anatomical normalisation; HB and PB performed the sleep analysis and classification; BH performed the analysis on disorders of consciousness patients. BH, LN and PB performed the statistical analysis. JS and JI provided inputs during the study. PB, BH and LN wrote the paper and the supplementals/method section. All authors discussed the data and provided feedbacks and suggestions.

---

**Figure figure supplement 1:**
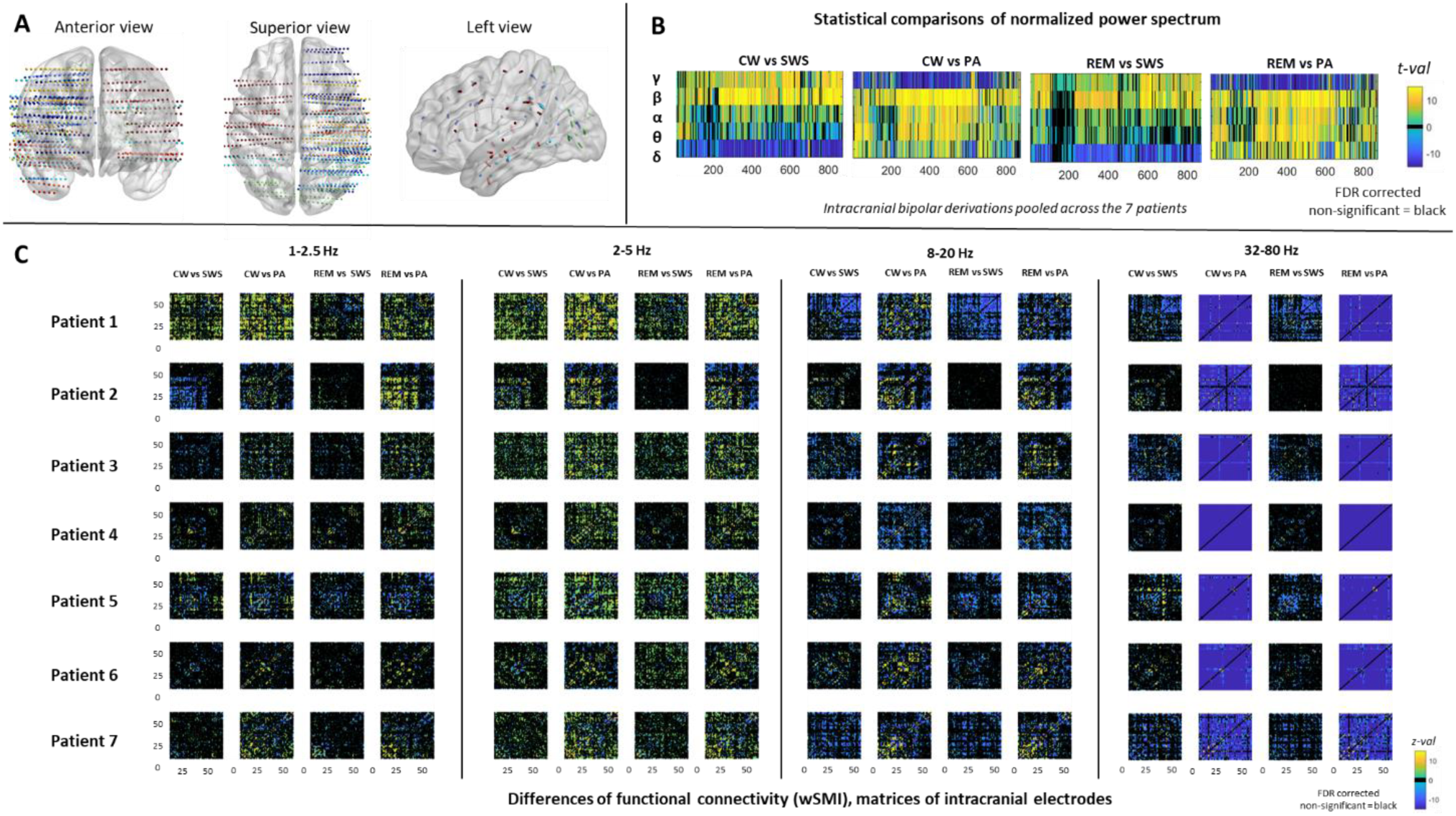
Individual anatomical data, power spectrum analysis and functional connectivity matrices. (a.) Representation in the MNI space of all electrodes implanted in the 7 patients recorded in the four distinct stages (including PA). Each patient is associated to a single colour. (b.) Power spectrum comparisons between conscious and unconscious conditions (CW/SWS, CS/PA, REM/SWS, REM/PA), computed for each bipolar contact in each of the classical 5 EEG frequency bands. The electrodes of the 7 patients are sorted according to their anterior-posterior coordinate (x axis). Only significant FDR-corrected t-values are plotted. Non-significant effects are represented in black. (c.) Matrices (bipolar derivation x bipolar derivation) showing the z-values of the statistical comparisons between each bipolar derivation across the different stages for three ranges of frequencies: 2-5 Hz; 8-20 Hz; 32-80 Hz. Only significant FDR-corrected z-values are represented. Non-significant effects are plotted in black. CW = conscious wakefulness; SWS = N3 Slow Wave Sleep; REM = Rapid Eye Movement Sleep; PA = propofol induced general anaesthesia

**Figure figure supplement 2:**
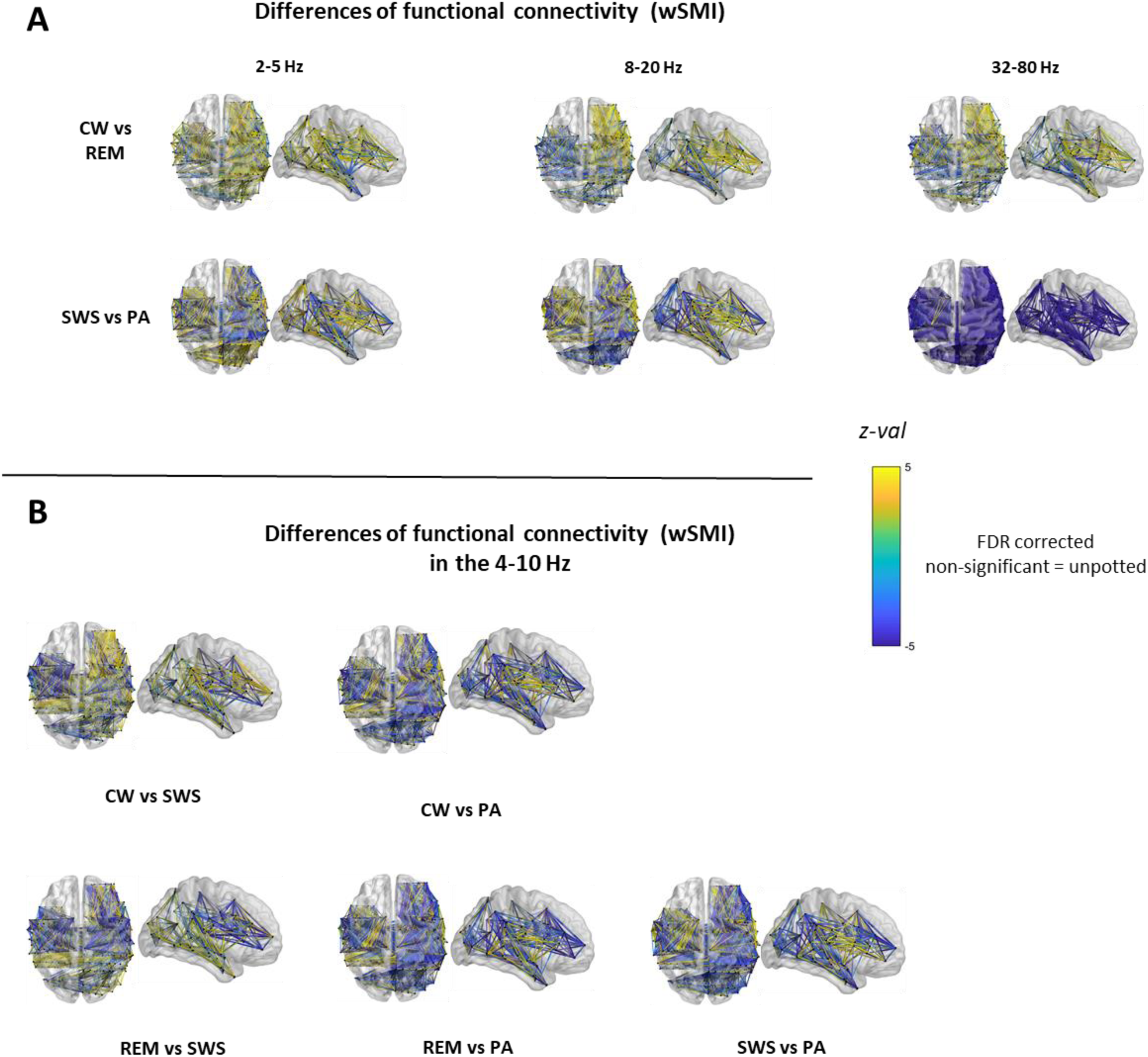
Additional functional connectivity data. (a.) Data from the 7 patients are plotted in the common MNI anatomical space. Comparisons of wSMI values that are absent in the main figure are displayed here: 2-5 Hz, 8-20 Hz and 32-80 Hz frequency bands. Only significant FDR-corrected z-values are represented (b.) Replication of the wSMI comparison display shown in main figure (panel B), adapted to the 4-10 Hz range. CW = conscious wakefulness; SWS = N3 Slow Wave Sleep; REM = Rapid Eye Movement Sleep; PA = propofol induced general anaesthesia

